# *S*. Typhimurium impairs glycolysis-mediated acidification of phagosomes to evade macrophage defense

**DOI:** 10.1101/2021.01.14.426635

**Authors:** Saray Gutiérrez, Julia Fischer, Raja Ganesan, Gökhan Cildir, Martina Wolke, Alberto Pessia, Peter Frommolt, Vincenzo Desiderio, Vidya R Velagapudi, Nirmal Robinson

## Abstract

Regulation of the cellular metabolism is now recognized as a crucial mechanism for the homeostasis of innate and adaptive immune cells upon diverse extracellular stimuli. Macrophages, for instance, increase glycolysis upon stimulation with pathogen-associated molecular patterns (PAMPs). Conceivably, pathogens also counteract these metabolic changes for their own survival in the host. However, despite this dynamic interplay in host-pathogen interactions, the role of immunometabolism in the context of intracellular bacterial infections is still unclear. Here, employing unbiased metabolomic and transcriptomic approaches, we investigated the role of metabolic adaptations of macrophages upon *Salmonella enterica* serovar Typhimurium (*S*. Typhimurium) infections. Importantly, our results suggested that *S*. Typhimurium abrogates glycolysis and its modulators such as insulin-signaling to impair macrophage defense. Mechanistically, glycolytic enzyme aldolase A is critical for v-ATPase assembly and the acidification of phagosomes upon *S*. Typhimurium infection, and impairment in the glycolytic machinery eventually leads to decreased bacterial clearance and antigen presentation in macrophages. Collectively, our results highlight a vital molecular link between metabolic adaptation and phagosome maturation in macrophages, which is targeted by *S*. Typhimurium to evade cell-autonomous defense.

## Introduction

Macrophages are sentinel immune cells playing pivotal roles in the host defense. They not only engulf and degrade the pathogens, but also secrete cytokines and present antigens to T cells to mount an effective adaptive immune response (1). Several pathogens such as *Salmonella enterica* serovar Typhimurium (*S*. Typhimurium) are restrained in phagosomes after being quickly phagocytosed by macrophages. However, *S*. Typhimurium also has evolved mechanisms to evade the hostile milieu of lysosomes and induce inflammatory cell death in macrophages. We have previously shown that *S*. Typhimurium induces type I interferon (IFN-I)-dependent and receptor-interacting serine/threonine-protein kinase 3 (RIP3)-mediated necroptosis in macrophages (2). It is also known that pro-inflammatory, necrotic cell death is associated with energy deficiency and metabolic instability in the cells (3). For instance, transfer of IFN-I receptor (IFNAR)-deficient or RIP3 kinase-deficient macrophages (that are cell death resistant) to wild type (WT) mice promotes better control of the pathogen implying that metabolically stable macrophages are more efficient in the control of pathogens (2).

A balanced immune response and metabolic homeostasis against invading pathogens are vital. Because substantial amount of energy is consumed when cells respond to immune stimuli, it is essential that they metabolically adapt to the demand (4). Recent studies highlighted the metabolic adjustments macrophages and dendritic cells undergo upon toll like receptor 4 (TLR4) activation with lipopolysaccharide (LPS) (5, 6). It has also been suggested that classically activated macrophages (M1), which respond readily to bacterial infections, derive their energy predominantly through glycolysis. On the other hand, alternatively activated macrophages (M2), which help in maintaining tissue homeostasis, obtain their energy mainly through oxidative phosphorylation (OXPHOS) (7, 8). Notably, metabolic intermediates arising from different metabolic pathways also significantly modulate the inflammatory response in immune cells. For instance, intracellular metabolites such as dimethyl fumarate (DMF) and itaconate have recently been found to modulate the innate and adaptive immune responses (9, 10). Similarly, tricarboxylic-acid (TCA) cycle intermediate succinate also modulates inflammation through Hypoxia-inducible factor 1-alpha (HIF-1α) in M1 macrophages (11). Thus, there is a dynamic crosstalk between metabolic intermediates and innate immune responses. Pathogens such as *S*. Typhimurium could also target this crosstalk and impair the metabolic homeostasis in macrophages. It has been reported that *S*. Typhimurium persists in M2 macrophages in a long-term infection model by sustaining fatty acid metabolism (12). Furthermore, *S*. Typhimurium also depend on its glycolysis machinery for survival in macrophages (13). We had also reported that the pathogen targets energy sensors such as AMPK and Sirtuin 1 for lysosomal degradation (14). More recently, we had shown that *S*. Typhimurium enhances leptin signaling to evade lysosomal degradation in macrophages (15). However, the implications of the metabolic pathways in macrophage defense against invading pathogens are largely unknown.

To understand the metabolic perturbations induced by *S*. Typhimurium, we performed an integrative metabolomics and transcriptomics analysis on macrophages infected with *S*. Typhimurium. This combined omics approach has revealed that glycolysis and its associated signaling pathways, such as insulin signaling facilitating glycolysis, are significantly down regulated upon *S*. Typhimurium infection. Furthermore, we show that down regulation of glycolysis by direct chemical inhibition or by genetically disrupting insulin-signaling in myeloid cells leads to elevated bacterial burden and impaired antigen presentation as a result of reduced acidification of phagosomes. Importantly, we also demonstrate that glycolysis regulates the assembly of vacuolar-type H^+^-ATPase complex (v-ATPase) and hence the acidification of phagosomes and glycolytic enzyme aldolase A critically regulates this process. Overall, our findings suggest that the Warburg-like-effect observed in macrophages upon infection is critical for the phagosome/lysosome-mediated clearance of pathogens. Moreover, pathogens such as *S*. Typhimurium have evolved strategies to disrupt this immunometabolic homeostasis in macrophages.

## Results

### *S*. Typhimurium infection promotes metabolic reprogramming in macrophages

To comprehensively characterize the metabolic alterations caused by *S*. Typhimurium infection, we carried out mass spectrometric analysis of metabolites in bone marrow-derived macrophages (BMDMs) infected with *S*. Typhimurium for 2h. As depicted in the PLSDA plot (**Figure S1A**) and in the heat map analysis of metabolites (**Figure 1A** for top-25 altered metabolites, **Figure S1B** for all metabolites analyzed), *S*. Typhimurium-infected macrophages presented a distinct metabolic profile when compared to uninfected (UI) controls. Metabolic pathway enrichment analysis revealed that carbohydrate-metabolism, which provides pyruvate for mitochondrial metabolism, and insulin signaling, which regulates glycolysis, were among the highly enriched pathway components upon *S*. Typhimurium infection (**Figure 1B**). Energy metabolites such as NAD+ was also highly down regulated upon *S*. Typhimurium infection (**Figures S1C**).

**Figure 1:**
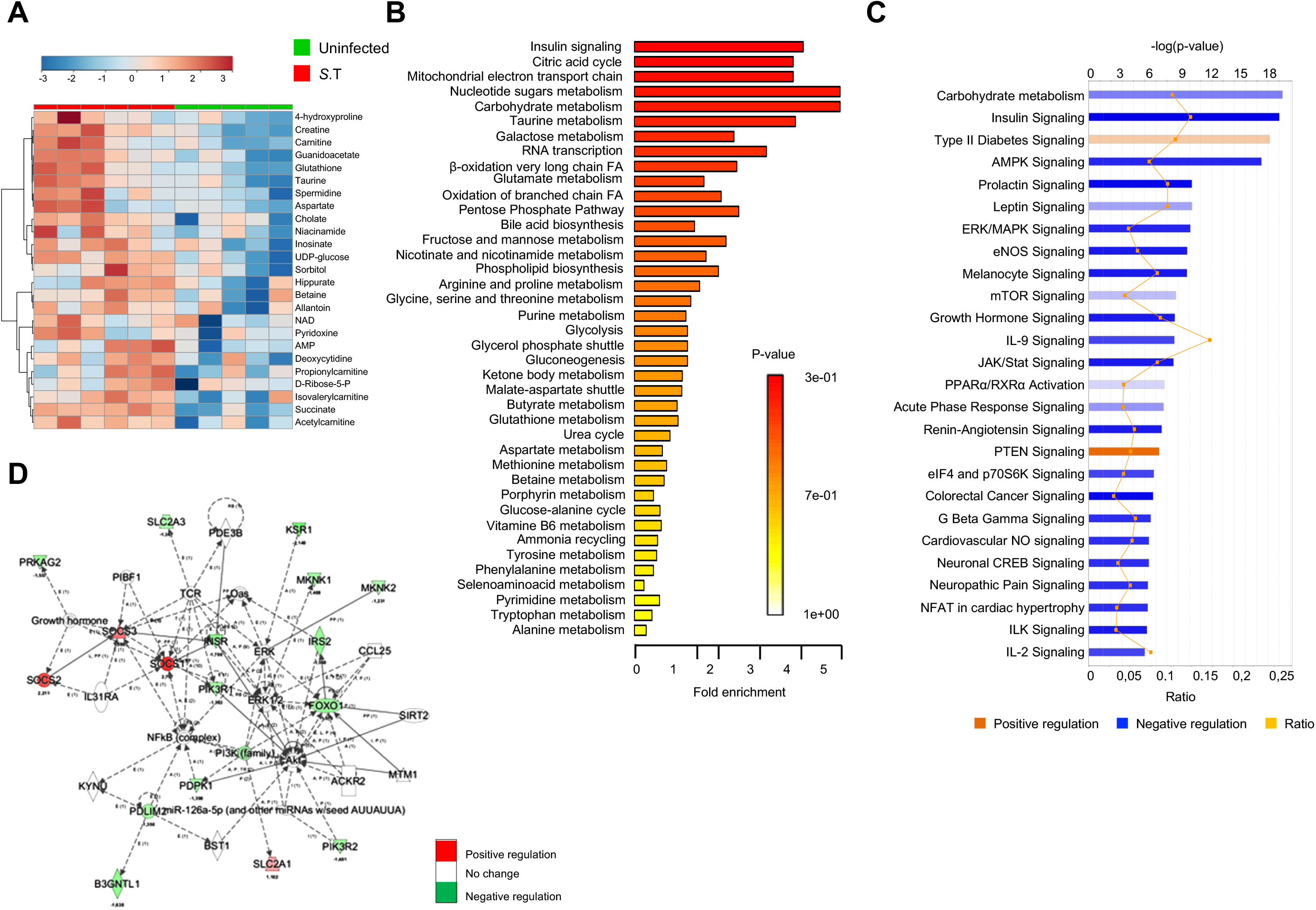
*S*. Typhimurium infection promotes metabolic reprogramming in macrophages. (**A**) Heatmap representation of 2-way hierarchical clustering of top-25 altered metabolites in BMDMs upon *S*. Typhimurium infection (2 h p.i.) (n=6) compared to uninfected controls (n=5). (**B**) Metabolic pathway enrichment analysis of metabolomics data from *S*. Typhimurium-infected BMDMs (2 h p.i) compared to uninfected controls. (**C**) Ingenuity pathway analysis of genes differentially expressed in RNA-seq data from *S*. Typhimurium-infected BMDMs (2h p.i) compared to uninfected controls (n=3). (**D**) Relative expression of genes from the insulin-signaling pathway in *S*. Typhimurium-infected BMDMs (2h p.i.) normalized to uninfected controls (n=3).

Complementary to this metabolite analysis, we also performed RNA-sequencing (RNA-seq) in BMDMs infected with *S*. Typhimurium at the same time point (2h). Consistent with the decrease in metabolites of glycolysis, RNA-seq data from BMDMs also showed that levels of genes involved in carbohydrate metabolism and insulin signaling were significantly down regulated (**Figures 1C, 1D** and **S1D**). Western blot analysis further confirmed that the expression levels of insulin receptor (IR) and its downstream target phosphorylated glycogen synthase kinase 3 (p-GSK3) were reduced upon *S*. Typhimurium infection **(Figure S1E and Figure S1F)**. In contrast, negative regulators of insulin signaling such as suppressors of cytokine signaling (SOCS) (16) and PTEN signaling pathway components (17) were up regulated (**Figures 1C and 1D**). Thus, metabolomics and transcriptomics together indicate that *S*. Typhimurium infection downregulates glycolysis and insulin-signaling that facilitates glycolysis.

### Virulence dependent inhibition of glycolysis in *S*. Typhimurium infected macrophages

Macrophages are known to undergo a switch in metabolism from OXPHOS to glycolysis upon various extracellular stimuli (18). However, the metabolic changes that occur upon intracellular bacterial infections are less understood. To show that glycolysis is indeed targeted by *S*. Typhimurium, we specifically analyzed the metabolites derived from glycolysis. This analysis confirmed that most of the metabolites generated upon breakdown of glucose were decreased, indicating that glucose flux was reduced upon infection with *S*. Typhimurium (**Figures 2A** and **S2A**). Consistently, *S*. Typhimurium infection of macrophages resulted in a decline in extracellular acidification rate (ECAR) at 4h, indicating reduced glycolytic flux, a phenomenon that was not observed upon LPS treatment (**Figure 2B**). Moreover, western blot analysis showed that the expression of Glut1, the main glucose transporter in macrophages (19), was significantly increased early upon infection (0.5-2h) followed by a decline during the later phase of infection (4h) (**Figure 2C**). In line with this, the levels of glucose-sensitive transcription factor MondoA and HIF-1α were also transiently up regulated (0.5-2h) and then down regulated over time (4h) (**Figure 2D and 2E**). Importantly, the change in the levels of Glut1 and the glucose-responsive transcription factors also correlated with the glucose uptake. Uptake of fluorescent glucose analogue 2-NBDG immediately upon infection was increased followed by a steady decline during the course of *S*. Typhimurium infection (**Figure 2F**). Our transcriptomics analysis revealed that majority of the glycolytic genes were down regulated upon *S*. Typhimurium infection in comparison to uninfected controls (**Figure S2B**). Despite an increase in glucose intake during the early phase of infection, the glycolytic metabolites were declined. Therefore, it is conceivable that *S*. Typhimurium actively blocks the proportionate up regulation of genes that are required to regulate the glycolytic flux.

**Figure 2:**
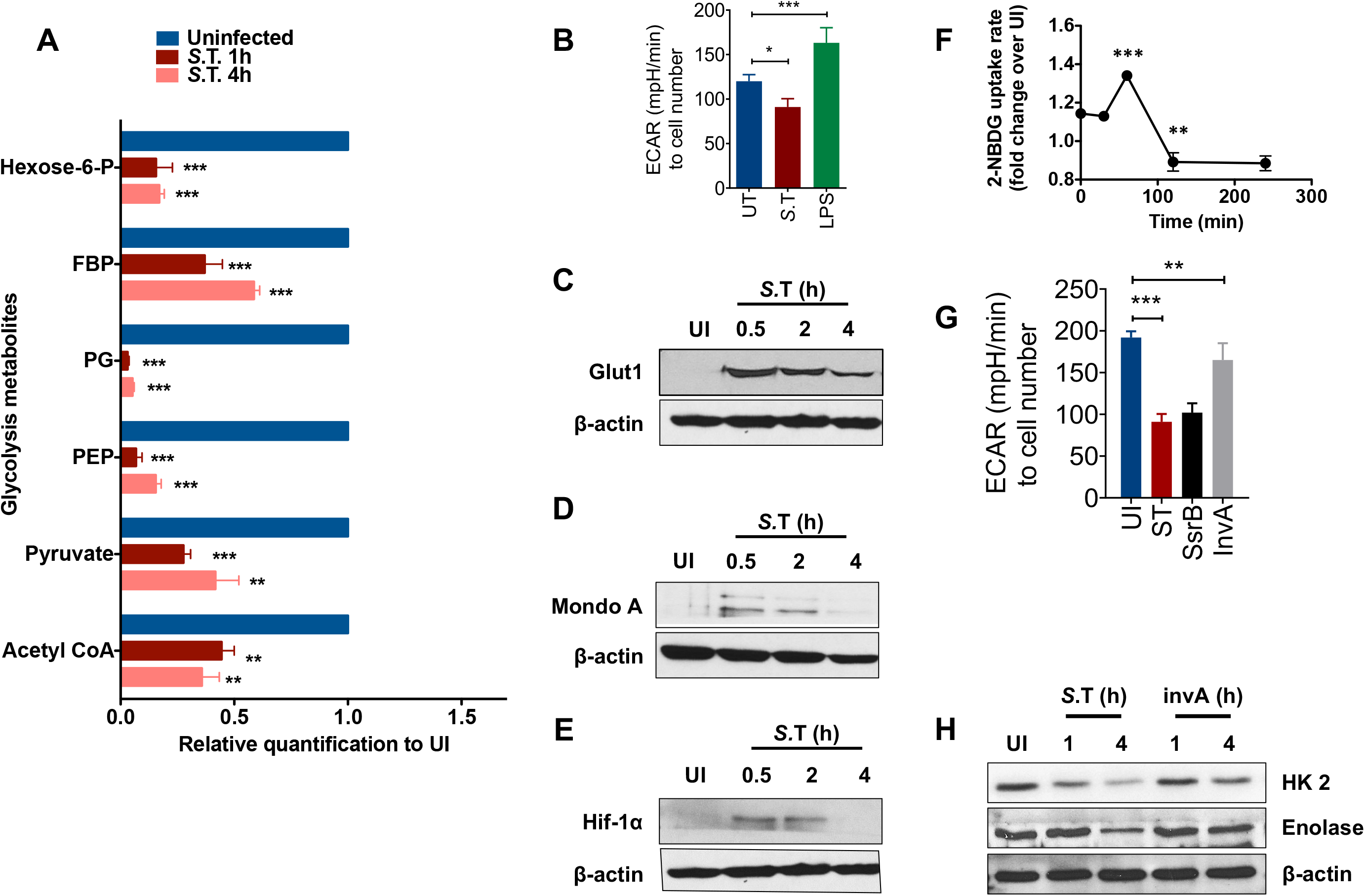
Virulence dependent inhibition of glycolysis in *S*. Typhimurium infected macrophages. (**A**) Abundance of glycolytic metabolites in *S*. Typhimurium-infected BMDMs after 1 h and 4 h post infection relative to uninfected (UI) BMDMs (n=6). (**B**) Extracellular Acidification Rate (ECAR) in BMDMs upon LPS treatment or S.T infection. Data is normalized to cell number (n=3). (**C**) Western blot analysis of Glut1 in *S*. Typhimurium-infected BMDMs compared to uninfected controls. β-actin was used as loading control. Image shown is representative of 4 independent experiments. (**D**) Immunoblot analysis of MondoA levels in different time points upon *S*. Typhimurium infection in BMDMs. (**E**) Immunoblot analysis of HIF-1a levels in different time points upon *S*. Typhimurium infection in BMDMs. (**F**) Kinetics of glucose-uptake (shown as 2-NBDG MFI) in *S*. Typhimurium-infected BMDMs relative to uninfected (UI) BMDMs analyzed by flow cytometry (n=3). (**G**) Extracellular Acidification Rate (ECAR) in BMDMs infected with WT *S*. Typhimurium, and *invA* and *ssrB mutants*. Data is normalized to cell number (n=6). (**H**) Immunoblot analysis of the protein levels of HK2, Glut1 and Enolase upon infection with WT *S*. Typhimurium or *invA* mutant at different time points. β-actin was used as a loading control. Data are shown as mean ± S.E.M. and statistical significance calculated using student t-test is represented as *=p<0.05; **= p<0.01; ***=p<0.001.

Next, we asked if the down regulation of glycolysis is a pathogenic mechanism of *S*. Typhimurium. Importantly, we observed that infection with heat-killed *S*. Typhimurium did not decrease the uptake of 2-NBDG (**Figure S2C**). To gain further insights into the virulence-dependent regulation of macrophage glycolysis by *S*. Typhimurium, we investigated the ability of two different *S*. Typhimurium mutants (known as *ssrB* and *invA*) to modulate the glycolytic response upon infection. Remarkably, *S*. Typhimurium mutant defective for expression of *invA*, a component of *Salmonella* pathogenicity island 1 (SPI-I), was significantly impaired in its ability to modulate glucose intake **(Figure S2D)** and ECAR **(Figure 2G)**. On the other hand, *S*. Typhimurium mutant defective for SPI-2-encoded transcriptional regulator *ssrB* was not impaired in its ability to regulate these parameters **(Figure S2D and 2G)**. Furthermore, the *invA* mutant *S*. Typhimurium was also unable to regulate the levels of Glut1 and glycolytic enzymes HK-2 and Enolase upon infection in macrophages **(Figure 2H)**. Overall, our analysis suggest that *S*. Typhimurium infection modulated glycolysis is virulence dependent.

### Macrophages depend on glycolysis for the clearance of intracellular bacteria

Since we found that *S*. Typhimurium downregulates glycolysis during the later phase of infection in BMDMs, we sought to determine the significance of glycolysis in the macrophage defense against *S*. Typhimurium. As the predominant function of macrophages is to eliminate invading pathogens, we studied the ability of BMDMs to degrade *S*. Typhimurium following a pre-treatment with the metabolically inactive glucose analogue 2-Deoxyglucose (2-DG) that inhibits glycolysis. Importantly, we found increased number of bacteria in macrophages when glycolysis was inhibited with 2-DG (**Figure 3A**). Similarly, inhibition of glucose uptake by Glut1 inhibitor Fasentin also increased the number of intracellular bacteria after 24h of infection (**Figure S3A**). It is well known that insulin signaling modulates glycolysis by regulating the cellular intake of glucose (20) and it has also been shown that myeloid-specific insulin receptor (IR) deficiency alters inflammation (21). We further show that glucose uptake (**Figure S3B**) was reduced in IR^Δmyel^ BMDMs compared to that of the IR^fl/fl^ (WT) and also observed a reduction in Glut1 (**Figure S3C**). Therefore, we next examined whether IR deficiency affects cell-autonomous defense against *S*. Typhimurium in macrophages. IR-deficient macrophages had increased bacterial burden upon infection with *S*. Typhimurium (**Figure 3B**). However, treatment of macrophages with recombinant insulin did not enhance the elimination of *S*. Typhimurium (**Figure S3D**). This is not surprising as our data show that insulin receptor and the downstream signaling are down regulated upon *S*. Typhimurium infection **(Figures 1D and S1F and S1G)**. Strikingly, when glucose uptake (**Figure S3E**) and ECAR (**Figure S3F**) were enhanced upon treatment with the glycolysis activator 4-hydroxytamoxifen (4-OHT) (22, 23), bacterial burden significantly decreased in macrophages (**Figure 3C**). Importantly, inhibition of glycolysis using 2-DG prevented enhanced bacterial clearance triggered by the treatment of 4-OHT (**Figure S3G**), confirming that 4-OHT-mediated enhanced bacterial elimination is glycolysis-dependent. Moreover, we found that the requirement of glycolysis for the clearance of intracellular bacteria is not specific for *S*. Typhimurium. Pathogens such as *Listeria monocytogenes (L. monocytogenes)* (**Figure 3D**) *and Staphylococcus aureus (S. aureus)* (**Figure 3E**) also survived better in 2-DG-treated macrophages. Similarly, IR-deficient macrophages also showed reduced ability to eliminate *S. aureus* (**Figure 3F**). Together, our results suggest that glycolysis plays a significant role in the elimination of intracellular bacteria.

**Figure 3:**
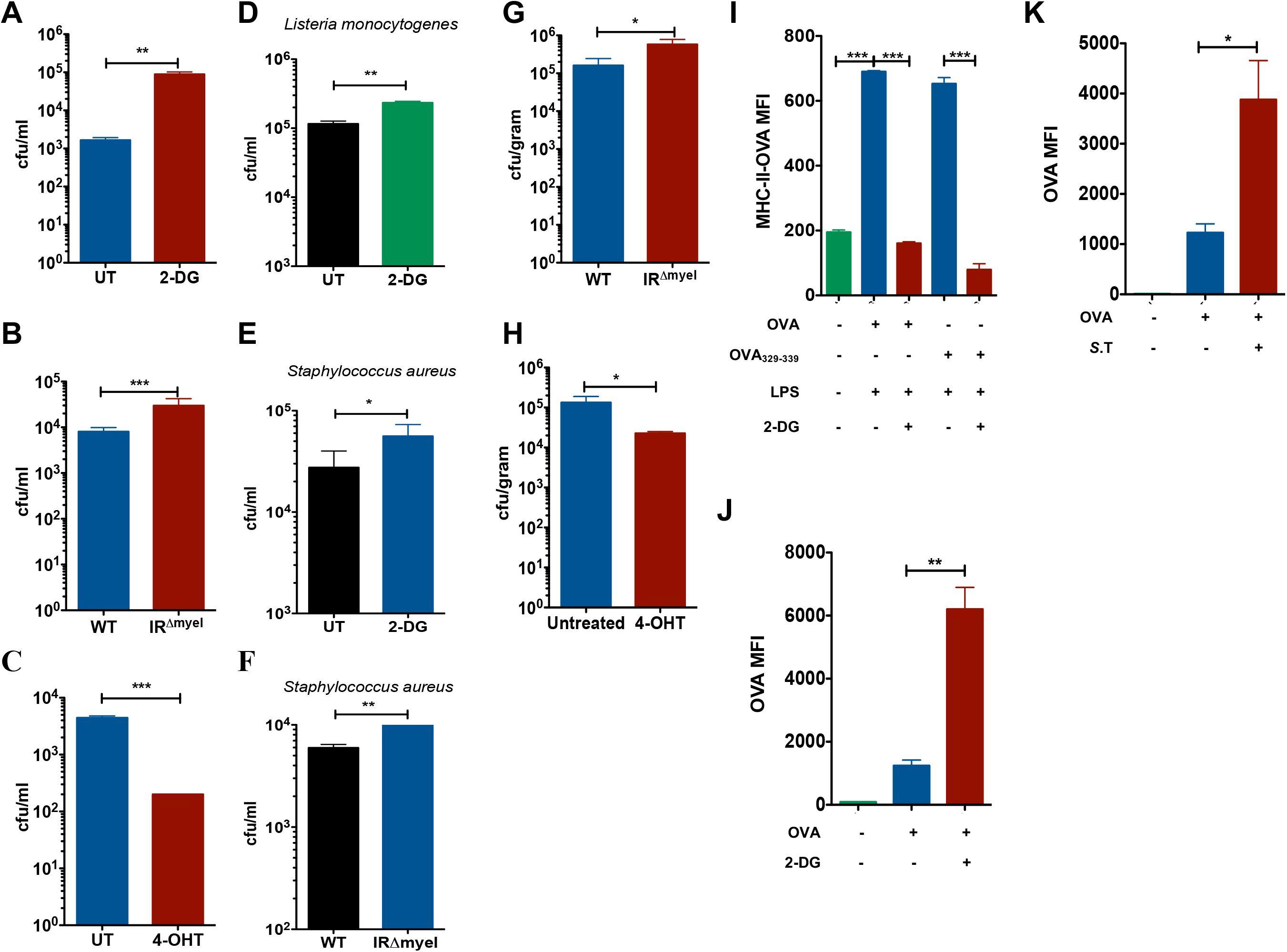
Macrophages depend on glycolysis for the clearance of intracellular bacteria. (**A**) Bacterial burden expressed as colony forming units (CFU) after 24h of *S*. Typhimurium infection in 2-DG-treated WT BMDMs compared to untreated (UT) controls (n=5). (**B**) Intracellular *S*. Typhimurium load in WT and IR^Δmyel^ BMDMs after 24h of infection (n=3). (**C**) *S*. Typhimurium in 4-OHT-treated WT BMDMs compared to untreated (UT) controls after 24h of infection (n=3). (**D**) *L. monocytogenes* burden in 2-DG-treated WT BMDMs compared to untreated (UT) controls (n=3). (**E**) *S. aureus* in 2-DG-treated WT BMDMs compared to untreated (UT) controls (n=3). (**F**) *S. aureus* in WT and IR^Δmyel^ BMDMs (n=3). (**G)** Bacterial load in livers of WT and IR^Δmyel^ mice after 3 days of *S*. Typhimurium infection. Data represents 2 experiments with 5 mice each. (**H**) Bacterial load in livers of 4-OHT-treated mice after 3 days of *S*. Typhimurium infection compared to untreated controls Data represents 2 experiments with 5 mice each. (**I**) MFI of OVA_323-339_-MHC II complexes on the surface of WT BMDMs pre-treated with 2-DG and LPS (n=3). (**J**) MFI of unprocessed Alexa647-labelled OVA in phagosomes isolated from 2-DG-treated BMDMs analyzed by flow cytometry (n=3). All samples were pre-stimulated with LPS. (**K**) MFI of unprocessed Alexa647-labelled OVA in bead containing phagosomes isolated from *S*. Typhimurium-infected BMDMs analyzed by flow cytometry (n=3). Data are shown as mean ± S.E.M. and statistical significance calculated using student t-test and represented as *=p<0.05; **= p<0.01; ***=p<0.001.

Consistent with the increase in bacterial burden, secretion of pro-inflammatory cytokines IL-6 and TNF-α was also increased when 2-DG-treated macrophages were infected with *S*. Typhimurium **(Figure S3H)**, *L. monocytogenes* (**Figure S3I**) or *S. aureus* (**Figure S3J**) or when IR^Δmyel^ macrophages were infected with *S*. Typhimurium (**Figure S3K**). In line with these findings, *S*. Typhimurium-induced IL-6 and TNF-α levels were decreased when glycolysis was induced with 4-OHT (**Figure S3L**). Increased cytokine secretion upon 2-DG treatment also correlated with the enhanced activation of NF-κB and p38 MAPK (**Figure S3M**). Next, we sought to investigate the involvement of glycolysis to control *S*. Typhimurium infection *in vivo*. Consistent with the *in vitro* results obtained in BMDMs, IR^Δmyel^ mice had increased bacterial burden in the liver after 3 days of *S*. Typhimurium infection compared to WT controls (**Figure 3G**). In contrast, 4-OHT-treated WT mice had significantly reduced *S*. Typhimurium in the liver (**Figure 3H**). Taken together, these results clearly suggest that increased glycolysis is beneficial for the clearance of bacteria *in vivo*.

Having found that glycolysis is required for the elimination of bacteria, we investigated whether the antigen processing and presentation could also be affected in macrophages when glycolysis is inhibited. To test this, we incubated 2-DG-treated and LPS-stimulated BMDMs with Ovalbumin (OVA) and analyzed the surface expression of the OVA peptide OVA_323-339_ bound to the MHC class II complex using specific antibodies by flow cytometry. We found that 2-DG-treatment drastically decreased the levels of OVA_323-339_-MHC II complexes on the surface of macrophages (**Figure 3I**). To gain better insight into the effect of glycolysis on antigen processing inside the phagosome, we incubated 2-DG-treated macrophages with beads coated with OVA conjugated to Alexa Fluor 647 dye. Analysis of the fluorescence intensity of the isolated phagosomes showed increased retention of Alexa647-OVA in 2-DG-treated macrophages suggesting reduced processing of the antigen (**Figure 3J**). Similarly, inhibition of glycolysis by *S*. Typhimurium infection also resulted in decreased processing of Alexa647-OVA as seen by the increased mean fluorescence intensity (MFI) in isolated Alexa647-OVA-coated-bead-containing phagosomes (**Figure 3K**). The decrease in processing of Alexa647-OVA caused by *S*. Typhimurium infection was partially rescued by treatment with 4-OHT (**Figure S3N**). Taken together, these data demonstrate that glycolysis is required for efficient antigen processing and antigen presentation.

### Glycolysis is crucial for phagosome maturation upon infection in macrophages

Macrophages engulf invading pathogens into phagosomes, which later fuse with lysosomes to degrade the pathogens. Increase in bacterial burden upon inhibition of glycolysis hinted that glycolysis could possibly regulate phagosomal functions in macrophages. To understand the role of glycolysis in phagosome maturation, we performed a series of flow cytometric assays to analyze the β-galactosidase and proteolytic activities in phagolysosomes containing inert beads in macrophages. To this end, beads either coated with C_12_FDG (a substrate for β-galactosidase) or with DQ-BSA (a substrate for proteases) were incubated with the macrophages. These substrates fluoresce when they react with their corresponding enzymes. Notably, 2-DG treatment prior to phagocytosis of beads showed markedly reduced β-galactosidase **(Figure 4A)** and proteolytic activities **(Figure 4B**) in bead-containing phagosomes. A similar decrease in the activities of β-galactosidase (**Figure 4C)** and proteases **(Figure 4D)** was also observed in IR-deficient macrophages when compared to WT controls. However, no significant differences in the phagocytosis of beads were observed in 2-DG-treated or IR-deficient macrophages compared to untreated or WT controls, respectively (**Figures S4A and S4B**). To test if *S*. Typhimurium mediated downregulation of glycolysis mimicked the effect of 2-DG, macrophages were infected for 2h to ensure that *S*. Typhimurium downregulated glycolysis. Cells were then allowed to phagocytose C_12_FDG-coated beads, which were chased into phagolysosomes and the activity of β-galactosidase was analyzed by flow cytometry. Interestingly, β-galactosidase activity on C_12_FDG-labelled beads phagocytosed after 2h of *S*. Typhimurium infection was reduced compared to the activity on C_12_FDG-labelled-beads phagocytosed by uninfected cells (**Figure 4E**). Furthermore, we found increased fluorescence signal from C_12_FDG-labelled beads that were phagocytosed after 30 min of *S*. Typhimurium infection, corresponding to the time when *S*. Typhimurium transiently increased glycolysis (**Figure S4C**). These results suggest that glycolysis is required for the efficient function of phagolysosomes and *S*. Typhimurium prevents this homeostasis during the later phase of infection by downregulating glycolysis.

**Figure 4:**
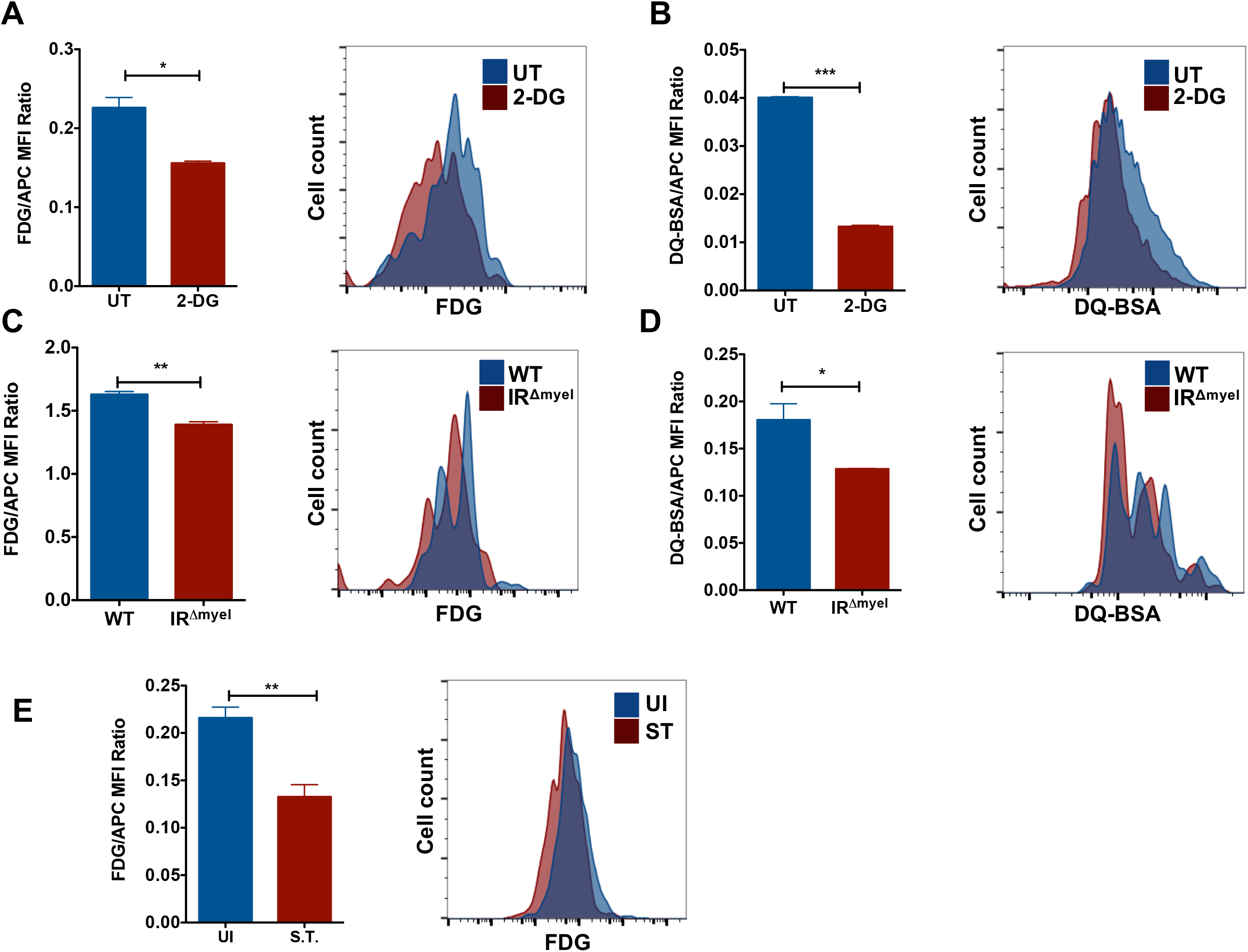
Glycolysis is essential for phagosome maturation. (**A**) Flow cytometry analysis of 2-DG pre-treated macrophages pulsed with C12FDG-coated beads (n=3) and (**B**) DQ-BSA-coated beads (n=3). Bar graphs represent mean fluorescence intensities (MFI) of C12FDG and DQ-BSA normalized to MFI of red fluorescence. MFI of (**C**) C12FDG and (**D**) DQ-BSA in WT and IR^Δmyel^ BMDMs normalized to MFI of red fluorescence (n=3). (**E**) Flow cytometric analysis of BMDMs infected with C12FDG and Alexa594-coated *S*. Typhimurium for 2h in WT BMDMs. Bar graphs represent mean MFI of C12FDG normalized to MFI of red fluorescence. Data are representative of at least three independent experiments with 3 replicates each. Data are shown as mean ± S.E.M. and statistical significance calculated using student t-test is represented as *=p<0.05; **= p<0.01; ***=p<0.001.

### Glycolysis critically regulates the acidification of phagosome and the assembly of v-ATPase complex

The activity of lysosomal enzymes and the maturation of the phagosomes to phagolysosomes are highly dependent on the acidification of the vesicle (24). Since inhibition of glycolysis impaired the activity of lysosomal enzymes in macrophages, we investigated if there is a defect in the acidification of the phagosomal lumen. Acidification of the phagosomes was studied using *E. coli* bioparticles labelled with pHrodo, a pH sensitive dye, which increases fluorescence intensity upon acidification. Notably, phagosome acidification was significantly reduced in bioparticle-containing phagosomes in macrophages pre-treated with 2-DG when compared to untreated controls (**Figure 5A**). Consistently, acidification was also limited in bioparticle-containing phagosomes of IR^Δmyel^ macrophages compared to WT controls (**Figure 5B**). Furthermore, *S*. Typhimurium-mediated inhibition of glycolysis also impaired acidification of bioparticle-containing phagosomes (**Figure 5C**), but the acidification was increased upon infection with *invA* mutant **(Figure S5A)**. Also, acidification of *S*. Typhimurium-containing phagosomes increased upon treatment with 4-OHT **(Figure S5A)**. Remarkably, the observed decrease in phagolysosome acidification in *S*. Typhimurium-infected macrophages was rescued when macrophages were treated with 4-OHT (**Figure 5D**), suggesting that *S*. Typhimurium prevents phagolysosome acidification by impairing glycolysis.

**Figure 5:**
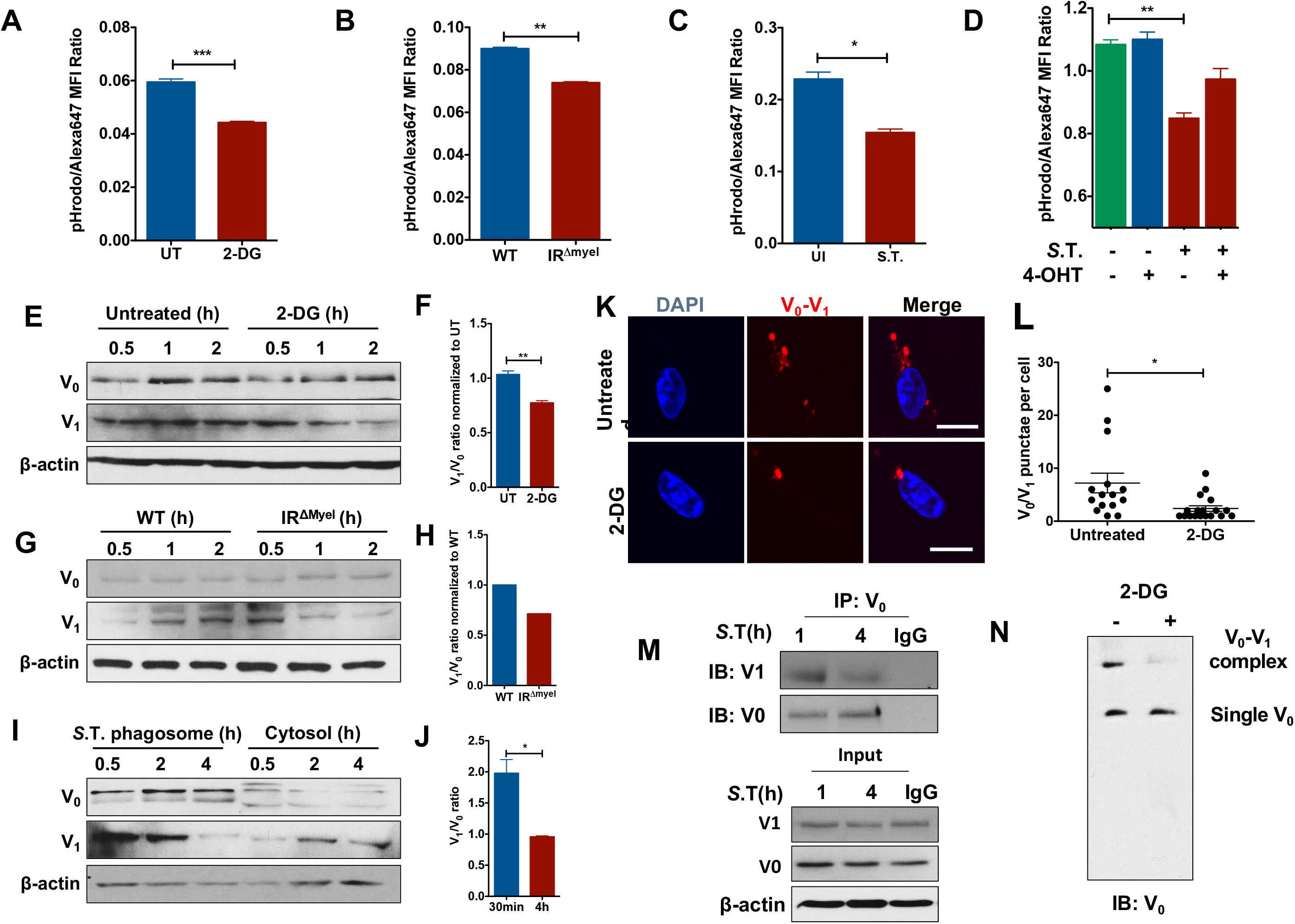
Glycolysis regulates phagosome acidification and v-ATPase assembly. (**A**) MFI of pH-sensitive pHrodo-*E. coli* particles in BMDMs untreated (UT) or pre-treated with 2-DG, normalized to Alexa647 MFI (n=3). (**B**) MFI of pHrodo-*E. coli* particles in WT and IR^Δmyel^ BMDMs normalized to Alexa647 MFI (n=3). (**C**) MFI of pHrodo-*E. coli* particles in *S*. Typhimurium-infected WT BMDMs (2h p.i.) normalized to Alexa647 MFI (n=3). (**D**) MFI of pHrodo-*E. coli* particles in 4-OHT-treated BMDMs infected with *S*. Typhimurium for 2h normalized to Alexa647 MFI (n=3). (**E**) Expression of v-ATPase subunits (V_0_, V_1_) in isolated bead-containing phagosomes from untreated and 2-DG-treated BMDMs. Image shown is representative of 3 individual experiments. (**F**) Immunoblot band intensities were quantified using imageJ and the V_1_/ V_0_ ratio was determined and plotted (n=3). (**G**) Expression of v-ATPase subunits (V_0_, V_1_) in isolated bead phagosomes from WT and IR^Δmyel^ BMDMs. (**H**) Western blot was quantified and V_1_/V_0_ ratios are shown. (**I**) Expression of v-ATPase subunits (V_0_, V_1_) in isolated *S*. Typhimurium phagosomes and cytoplasm. (**J**) V_1_/V_0_ ratios were quantified and plotted. **(K)** PLA analysis of v-ATPase subunits V_0_ and V_1_ interaction in 2-DG-treated BMDMs using confocal microscopy (scale bars indicate 10 µm). Image shown is representative of 3 individual experiments. (**L**) Quantification of V0 and V1 interaction in 2-DG treated BMDMs (n=15). (**M**) V_0_ subunit *a* was immunoprecipitated from isolated *S*. Typhimurium phagosomes and probed for V_1_ subunit B. (**N**) Phagosomes isolated from 2DG-treated and untreated macrophages were subjected to Native-PAGE and immunoblotted for v-ATPase subunits V_0_ and V_1_.

Acidification of phagosomes is mediated by a multimeric protein complex known as vacuolar-ATPase (v-ATPase), which is composed of 14 subunits organized in two main catalytic macro domains: V_0_ and V_1_ (25). While V_0_ is permanently bound to the membrane of phagosomes, V_1_ is located in the cytosol and gets actively recruited onto the phagosome to interact with V_0_ and thus activate the proton pump (26). To test whether inhibition of glycolysis could have an effect on the assembly of the v-ATPase complex in macrophages, isolated bead containing-phagosomes from 2-DG-treated BMDMs and IR^Δmyel^ BMDMs were analyzed for the expression of subunit-*a* and subunit-B which are part of the V_0_ and the V_1_ macro domains, respectively. V_0_ subunit-*a* was detected in comparable amounts in bead containing-phagosomes isolated from 2-DG-treated and untreated controls (**Figure 5E**). Similarly, abundance of V_0_ subunit-*a* was comparable in phagosomes isolated from IR^Δmyel^ macrophages and the WT controls (**Figure 5G**). However, the expression of the V_1_ subunit-B was reduced in the phagosomes isolated from 2-DG-treated macrophages (**Figure 5E and 5F**). Similarly, we found reduced levels of the V_1_ subunit-B in bead containing-phagosomes isolated from IR^Δmyel^ macrophages **(Figure 5G and 5H)** and from fasentin-treated macrophages when compared to WT controls (**Figure S5B**). Notably, we did not observe differential amounts of V_0_ and V_1_ subunits in the total cell lysates of 2-DG-treated macrophages compared to controls (**Figure S5C**) or in fasentin-treated cells compared to untreated controls (**Figure S5D**). These findings suggest that impaired glycolysis prevents the assembly of the v-ATPase complex rather than the expression itself. V_1_ recruitment on to phagosomes containing *S*. Typhimurium was also reduced, while V_0_ levels did not vary significantly between different time points (**Figures 5I**). The decline in V_1_ on phagosomes harboring *S*. Typhimurium corresponded with the time when glycolysis was inhibited (**Figure 2A-2E**). Similarly, Proximity Ligation Assay (PLA) also showed reduced interaction of V_0_ and V_1_ in 2-DG-treated macrophages (**Figures 5K and 5L**). The decreased interaction between V_0_ and V_1_ upon *S*. Typhimurium infection was also confirmed by PLA (**Figures S5E and S5F**). We also immunoprecipitated (IP) V_0_ from isolated *S*. Typhimurium-containing phagosomes and observed the complex formation of V_0_ and V_1_ however, V_1_ binding to V_0_ was reduced in *S*. Typhimurium-phagosomes isolated after 6h (**Figure 5M**). To further investigate the effect of the inhibition of glycolysis on v-ATPase complex formation, we conducted Native SDS-PAGE using isolated bead-containing phagosomes from macrophages treated or untreated with 2-DG. 2-DG treatment resulted in a significant decrease in the formation of v-ATPase complex (**Figure 5N**). Taken together, these results strongly suggest that glycolysis plays a critical role in the assembly of v-ATPase.

### Aldolase A critically regulates the assembly of v-ATPase and phagosome acidification

Glycolytic enzymes aldolase A and phosphofructokinase-1 (PFK1) have been shown to interact with different subunits of the v-ATPase in yeast, likely acting as scaffold proteins and are required for the acidification of endosomes (27, 28). Confocal microscopy confirmed that aldolase A colocalized with inert E-coli bioparticles-containing phagosomes when glucose was abundant **(Figures 6A)**. However, co-localization of aldolase A with *E. coli* particles-containing phagosomes was significantly reduced upon glycolysis-inhibition with 2-DG **(Figures 6A and 6B)**. Aldolase A colocalized with *S*. Typhimurium-containing phagosomes as early as 30 min post infection, but the amount of *S*. Typhimurium-phagosomes positive for aldolase A was reduced 4h post infection **(Figure 6C)**. PLA analysis revealed interaction between V_0_ and Aldolase A in untreated macrophages. However, the number of red puncta (indicating interaction between V_0_ and Aldolase A) was significantly reduced upon treatment with 2-DG **(Figure 6D)**. V_0_ and Aldolase A interaction was also observed in *S*. Typhimurium-infected macrophages but the frequency of puncta per cell reduced after 4h of infection compared to 30 min **(Figures 6E and 6F)**. Total levels of aldolase A also showed a modest increase in macrophages upon *S*. Typhimurium infection **(Figure 6G)**. Since we observed that the reduction in the recruitment of aldolase A on to bead or *S*. Typhimurium-containing phagolysosomes correlated with the decrease in phagosome acidification, we sought to determine if aldolase A played a role in the regulation of phagolysosome acidification. Short interfering RNA (siRNA)-mediated knockdown (KD) of Aldolase A in BMDMs **(Figure 6H)** significantly reduced phagosomal acidification as indicated by reduced pHrodo fluorescence in aldolase A-depleted BMDMs **(Figure 6I)**. As a direct consequence of reduced phagosome acidification, Aldolase A depletion also significantly inhibited phagosomal processing as evident from increased number of intracellular *S*. Typhimurium after 24h of infection **(Figure 6J)**. Taken together, our data signifies the roles of glycolysis and glycolytic enzyme aldolase A in the assembly of v-ATPase, vacuolar acidification and clearance of intracellular bacteria **(Figure 7)**.

**Figure 6:**
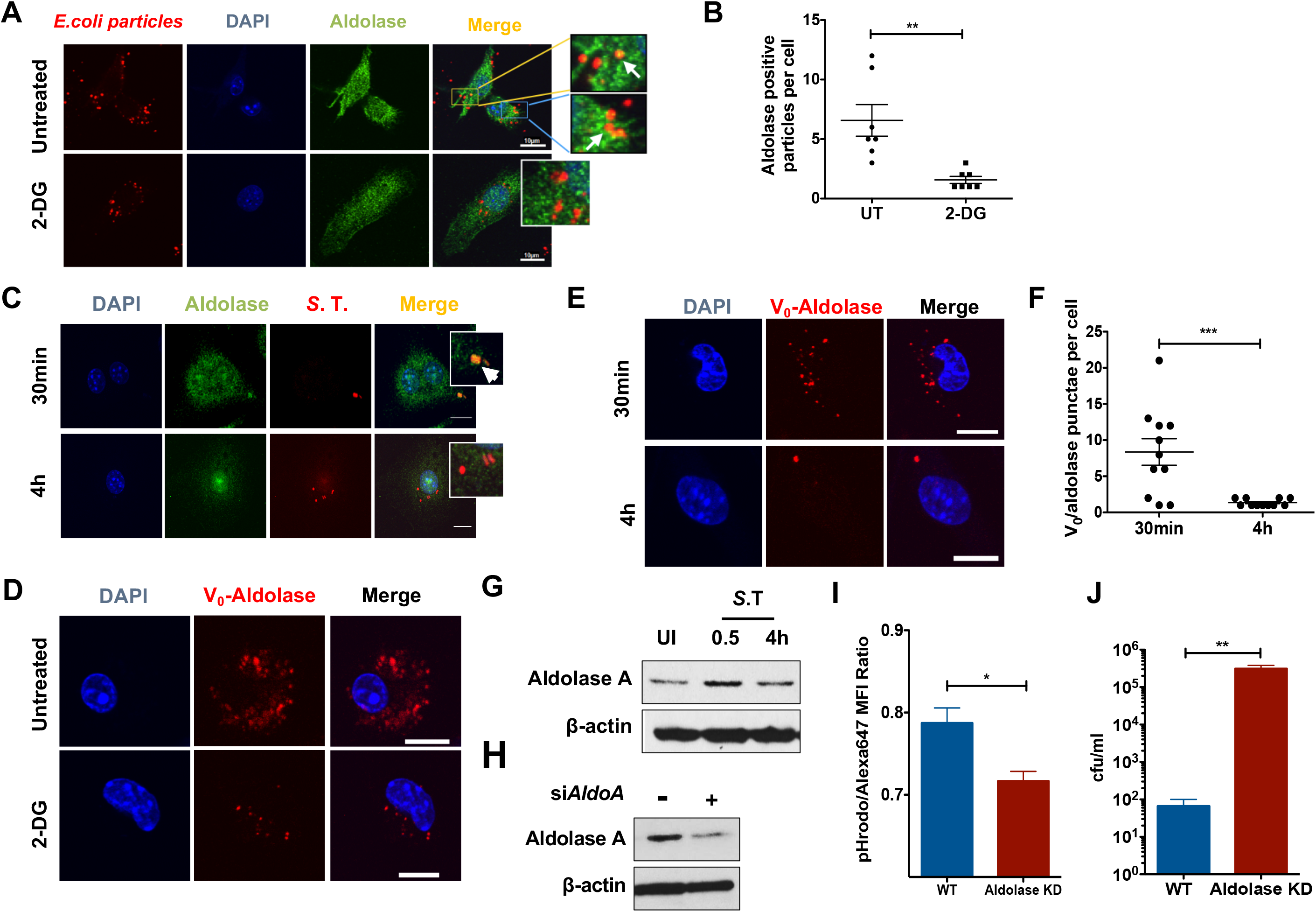
Aldolase A critically regulates the assembly of v-ATPase and phagosome acidification. (**A**) Confocal microscopy of 2-DG-treated BMDMs pulsed with E. coli inert fluorescent particles immunostained for aldolase A (green) (scale bars indicate 10µm). Image shown is representative of 3 individual experiments. **(B)** Quantification of aldolase A co-localization with E. coli in 2-DG-treated BMDMs (n=7). (**C**) Confocal microscopy of *S*. Typhimurium-infected (S.T., red) BMDMs immunostained for aldolase A (green). Image shown is representative of 3 individual experiments. **(D)** PLA analysis of V0-aldolase A interaction in 2-DG treated BMDMs. Image shown is representative of 3 individual experiments. **(E)** PLA analysis of V_0_-aldolase A interaction in *S*. Typhimurium-infected BMDMs. Image shown is representative of 3 individual experiments. **(F)** Quantification of V_0_ and aldolase A interaction in S. Typhimurium-infected BMDMs (n=10). **(G)** Western blot analysis of Aldolase A expression in BMDMDs infected with *S*. Typhimurium at indicated time points. **(H)** Knockdown of *aldolase A* in BMDMs using control siRNA and *aldolase A*-specific siRNA. **(I)** MFI of pHrodo-E. coli particles in aldolase A KD BMDMs normalized to Alexa647 MFI analyzed by flow cytometry (n=3). **(J)** Quantification of *S*. Typhimurium cfu in *aldolase A* KD macrophages 24h post-infection compared to WT controls (n=3).

**Figure 7:**
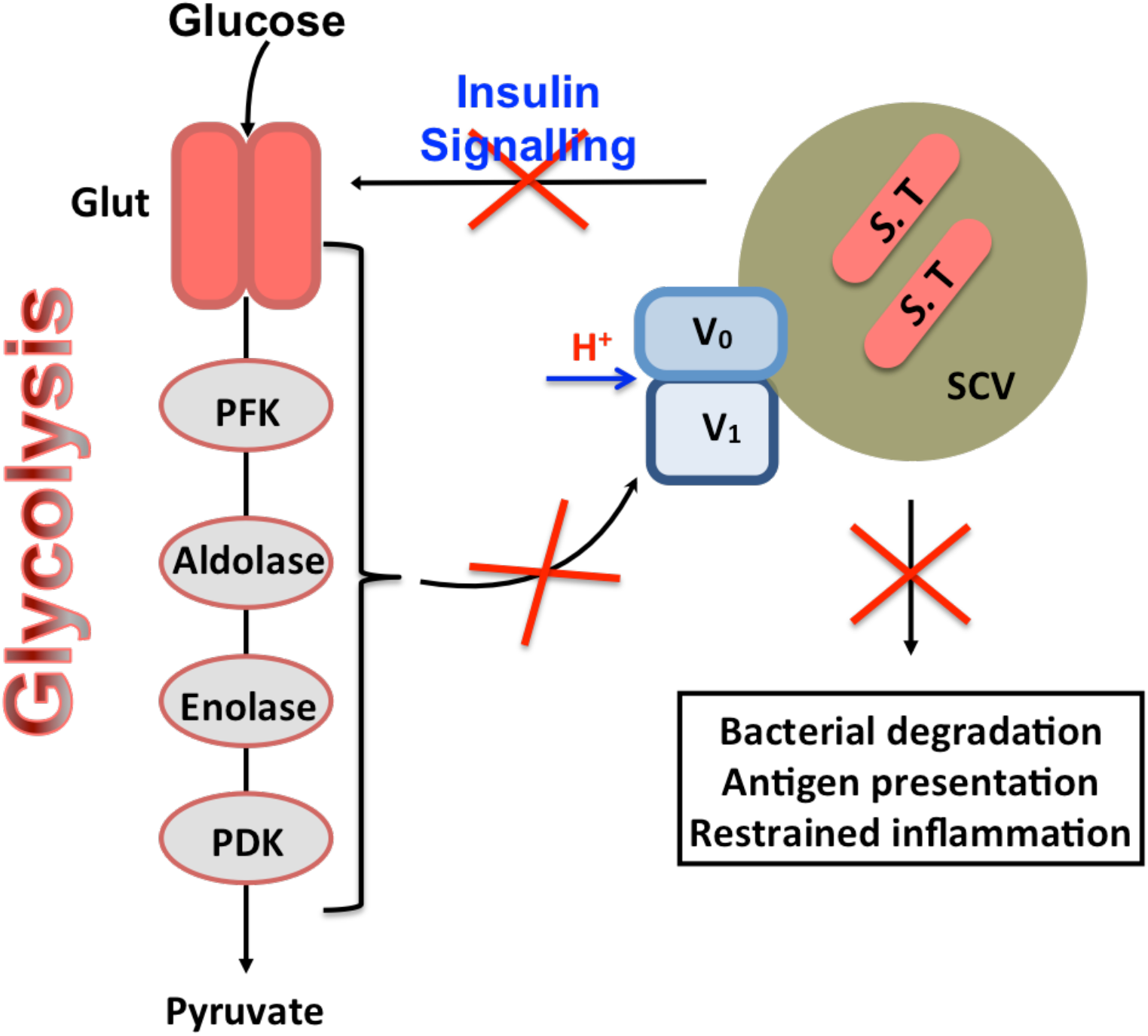
Aldolase A critically regulates the assembly of v-ATPase and phagosome acidification. Schematic representation of *S*. Typhimurium-mediated evasion of phagosome degradation in macrophages by preventing glycolysis-regulated assembly of the v-ATPase.

## Discussion

Upon pro-inflammatory stimuli, macrophages undergo metabolic reprogramming as part of an innate immune response. In this study, we deciphered that *S*. Typhimurium down regulates glycolysis, which is critical for phagolysosomal function and antigen-presentation in macrophages. We further demonstrate that glycolysis is required for the assembly of v-ATPase complex on phagosomes and acidification of phagosomes, which is coordinated by aldolase A.

An orchestrated and effective immune response requires high levels of ATP and biomolecules. However, sites of infection are often poor in oxygen and nutrients. It has become evident that, macrophages and dendritic cells undergo a metabolic shift similar to the Warburg-effect observed in cancer cells when pattern recognition receptors are activated to meet the high-energy demand (29). Similarly, intracellular pathogens have to adapt in order to survive in the hostile intracellular milieu, which requires altering the metabolic environment to its advantage. In this regard, it is interesting to note that pathogenicity of *Mycobacterium tuberculosis* has been linked to its ability to modulate cellular metabolism (30). *S*. Typhimurium is a facultative intracellular pathogen that is known to alter mitochondrial metabolism in host cells and reduce ATP production (14, 31-33). However, the specific regulation of host metabolic pathways and its interplay with innate immune mechanisms upon *S*. Typhimurium infection remain unknown. Our metabolomics and transcriptomics analysis converged in revealing that the central-carbon-metabolism pathways contributing to ATP generation in macrophages, namely glycolysis, the TCA cycle and the mitochondrial electron transport chain, were markedly down regulated upon infection with *S*. Typhimurium. Our results are in agreement with previous microarray analysis conducted on colon mucosa and liver of *S*. Typhimurium-infected mice, where a decline in OXPHOS and carbohydrate metabolism together with decreased levels of hormones regulating the metabolic pathways have been reported (34, 35). Furthermore, down regulation of these metabolic pathways is consistent with the previously reported necrotic cell death and ATP depletion induced by *S*. Typhimurium (36). Additionally, *S*. Typhimurium-mediated down regulation of glycolytic metabolites could be a direct effect of bacterial metabolism, since *S*. Typhimurium in macrophages has been shown to rely on glycolysis for its replication, reducing the availability of glucose in the host cytosol (13). Taken together, our results highlight glycolysis as a key target of *S*. Typhimurium during its interplay with host cells.

Emerging evidence suggests that Warburg effect-like metabolic shift observed in macrophages fuel inflammatory responses when exposed to TLR agonists (37). The metabolic shift towards glycolytic flux fuels both the TCA cycle and the pentose phosphate pathway (PPP) thus providing a double impact on the innate immune defense mechanisms. On the one hand, increased production of NADPH by the PPP results in ROS-mediated inflammation as a consequence of the transfer of electrons from NADPH to NADPH-oxidase (38). Metabolites such as succinate derived from TCA cycle have also been shown to induce IL-1β in a HIF-1α-dependent manner in macrophages stimulated with LPS (11). In contrast, our data indicate that *S*. Typhimurium-induced inflammation is independent of these metabolic mechanisms since levels of ribose-5-phosphate (PPP), and succinate (TCA cycle) declined. Moreover, inhibition of glycolysis further enhanced inflammatory cytokine secretion in *S*. Typhimurium-infected macrophages. Our data corroborates with the report that *S*. Typhimurium disrupts glycolysis to induce inflammasome-mediated inflammation (39). Therefore, we propose that *S*. Typhimurium inhibits glycolysis to evade phagosomal clearance leading to the activation of canonical inflammatory pathways.

A predominant function of macrophages is to phagocytose invading pathogens and eliminate them in phagolysosomes. Previous reports have shown that *S*. Typhimurium prevents fusion of lysosomes with bacteria-containing phagosomes (40). Contradictorily, several other studies have shown that *S*. Typhimurium-containing vacuoles (SCVs) are accessible to lysosomal markers and they fuse with lysosomes (41),(42). This paradox could be clarified by our findings, which show that macrophages initially upregulate glycolytic machinery to enhance acidification of the phagosomes and acquire late endosomal properties. However, *S*. Typhimurium uses the phagosomal acidic environment to express its SPI-2-encoded virulence factors (43) and inhibit phagosome maturation. This is evident from a rapid increase in Glut1 expression and glucose import followed by a marked reduction at the later phase of infection. Interestingly, we also observed that the reduction in glucose import and acidification of phagosomes is virulence associate and is partially SPI-I-dependent as the *S*. Typhimurium mutant *invA* did not abrogate glycolysis as that of the WT bacteria (**Figures 2G-2H**). Our findings clearly demonstrate that reduced glycolysis due to deficient insulin signaling or feeding cells with 2-DG leads to increased bacterial replication *in vitro* and *in vivo*. Accordingly, up regulation of glycolysis using 4-OHT starkly reduced bacterial burden in BMDMs and *in vivo*. These results indicate that metabolism is intricately linked to phagosomal functions in macrophages. Our investigations using inert biological particles and beads coated with substrates for various lysosomal enzymes reveal a general cellular mechanism that glycolysis regulates the activity of lysosomal enzymes. *S*. Typhimurium-mediated inhibition of glycolysis also reduced the processing and presentation of OVA peptides on MHC molecules. This is consistent with our previous report that *S*. Typhimurium-infected antigen presenting cells lack antigen presentation ability (44). Interestingly, both the processing of antigens and the activation of MHC II (cleavage of the invariant chain) are pH-dependent (45).

Acidification of the phagosomes is a prerequisite for the degradative function, because lysosomal enzymes require acidic pH for their optimal activity. Acidification is also required for the interaction with endocytic vesicles and the maturation of phagosomes themselves (46). Metabolic intermediates such as ROS catalyzed by NADPH-oxidase and generated in mitochondria juxtaposed to pathogen containing phagosomes (47) and NADH-dependent NO production by iNOS (48) have been shown to facilitate the bactericidal activity of phagolysosomes. Production of these metabolic intermediates also requires protons generated by v-ATPase complex. Acidification of phagosomes is regulated by the multimeric complex v-ATPase and is critical for the maturation of phagosomes, fusion of lysosome with phagosomes and the activity of lysosomal enzymes (49, 50). Since we observed defects in phagosome maturation in glycolysis-deficient macrophages, we reasoned that glycolysis could be involved in the acidification of phagosomes. As expected, under conditions of reduced glycolytic flux, acidification of phagosomes containing inert biological particles was significantly reduced, which is a result of defective assembly of V_0_ and V_1_ on phagosomes. Our study thus suggests that *S*. Typhimurium mediated down regulation of glycolysis prevents acidification of the pathogen-containing phagosomes by hampering the assembly of V_0_ and V_1_ on the phagosomes. This is consistent with a previous report demonstrating that high glucose availability facilitates the assembly of v-ATPase thus enabling increased entry of influenza virus (51). However, the mechanism remains poorly characterized.

We found that glycolytic enzymes were downregulated when glucose intake was reduced by *S*. Typhimurium infection, indicating that the import of glucose could be an additional mechanism regulating the expression of glycolytic enzymes. Consistently, we observed a decrease in MondoA over time, a glucose-induced transcription factor responsible for the transcriptional regulation of enzymes involved in carbohydrate catabolism, including glycolysis and PPP (52). Glycolytic enzymes are also known to perform multifaceted non-glycolytic functions such as transcriptional regulation (53), cell motility and regulation of apoptosis (54). Glycolytic enzymes such as aldolase and PFK have been directly implicated in the assembly of v-ATPase (27, 28). Future studies on the specific roles of different glycolytic enzymes in the regulation of phagosomes will be valuable in understanding the process of phagosome maturation and will expand its role in linking innate and adaptive immunity.

Taken together, our findings convincingly demonstrate that glycolysis critically regulates the phagolysosomal activity of macrophages, which is evaded by *S*. Typhimurium. Conceivably, in pathological conditions such as diabetes wherein cells are insensitive to insulin, patients become increasingly susceptible to infections because macrophages will be inefficient in eliminating pathogens as a result of reduced glycolysis. Furthermore, the significance of glycolysis in antigen processing and presentation could be applied in designing adjuvants for vaccines. Finally, increase in antimicrobial properties of macrophages upon 4-OHT treatment suggests an alternate approach to control drug-resistant pathogens.

## Experimental Procedures

### Ethics statement

All animal procedures were in accordance with institutional guidelines on animal welfare and were approved by the North Rhine-Westphalian State Agency for Nature, Environment, and Consumer Protection [Landesamt für Natur, Umwelt and Verbraucherschutz (LANUV) Nordrhein-Westfalen; File no: 84-02.05.40.14.082 and 84-02.04.2015.A443] and the University of Cologne.

### Preparation of bacteria

Bacteria from a single colony was inoculated into 5 mL of BHI medium and incubated overnight at 37°C with constant agitation. Next, 1 mL of bacteria suspension was transferred into 50 mL of BHI and grown until OD_600_:1 (logarithmic growth phase). Concentration of bacteria was estimated by plating serial dilutions on BHI agar plates. When indicated, *S*. Typhimurium was inactivated for 10 min at 95° C.

### Cell culture and bacterial infection

Bone marrow was extracted from femur of 8-12 weeks old female wild-type (WT) C57BL/6 mice (Charles River Laboratories) or insulin receptor knockout (IR^Δmyel^) mice (provided by Jens Brüning, MPI for Metabolism and Ageing, Cologne, Germany). Bone marrow was differentiated into macrophages for 7 days in RPMI medium supplemented with 20% L929 supernatant and 10% fetal bovine serum (FBS). Bone marrow derived macrophages (BMDMs) were infected with *Salmonella* Typhimurium SL1344 or *S. aureus or L. monocytogenes* at a multiplicity of infection (MOI) of 10. After infection, BMDMs were incubated with the bacteria for 10 min at RT and 30 min at 37°C. This incubation time was sufficient for bacteria to be internalized by macrophages as confirmed by microscopy analysis. Cells were then washed with medium containing 50 µg/ml of gentamicin and incubated in medium with 50 µg/ml of gentamicin. After 2h, the concentration of gentamycin was reduced to 10 µg/ml. BMDMs were treated with 1 mM 2-DG (Life Technologies) dissolved in medium without glucose for 2h prior to infection, 1 µM 4-hydroxytamoxifen (4-OHT; Sigma) for 1h at 37°C prior to infection and during the course of infection, oligomycin (Sigma) 2 µg/ml for 30 min prior to infection, pyruvate 1mM for 2h prior to infection or with *S*. Typhimurium LPS (Sigma) 100 ng/ml for the indicated times.

### Global metabolomics analysis

1×10^6^ cells per sample were taken for metabolomics analysis. Trypsinized cells were washed twice with PBS buffer and then with deionized water for few seconds. Subsequently, cells were quickly quenched in liquid nitrogen and stored at −80°C until further analysis. Frozen cell samples were thawed step wise at −20°C and 4°C and then metabolites were extracted by adding 20 μl of labeled internal standard mix and 1 ml of cold extraction solvent (90/10 ACN/H2O + 1% FA). Cells were then vortexed for 30sec, sonicated for 30 sec in three cycles, and incubated on ice for 10 min. After the centrifugation at 14000 rpm for 15 min at 4°C, 800 µl of supernatants were aspirated into Eppendorf tubes. The collected extracts were dispensed in OstroTM 96-well plate (Waters Corporation, Milford, USA) and filtered by applying vacuum at a delta pressure of 300-400mbar for 2.5 min on robot’s vacuum station. The clean extract was collected to a 96-well collection plate, placed under OstroTM plate. The collection plate was sealed and centrifuged for 15 min, 4000 rpm, 4°C and placed in auto-sampler of the liquid chromatography system for the injection. Isotopically labeled internal standards were obtained from Cambridge Isotope Laboratory. Inc., USA (Table S1). Instrument parameters, analytical conditions and data analysis were performed as previously described (55). Metabolomics data analysis was carried out using a web-based comprehensive metabolomics data processing tool, MetaboAnalyst 3.0 (56). Log-transformed and auto scaled data i.e., mean-centered and divided by the standard deviation of each variable, was used for various data analysis. t-test for unequal variances (Welch’s t-test) was applied by default to every compound.

### RNA-sequencing

Total RNA from *S*. Typhimurium-infected BMDMs was extracted 2 h post infection using the Qiagen RNAeasy kit and cDNA was synthesized with SuperScript III (Life Technologies) following the manufacturer’s instructions. For Illumina sequencing, libraries were prepared from total RNA with Ribo-Zero treatment according to the manufacturer’s instructions. The analysis was carried out using the standardized RNA-Seq workflow on the QuickNGS platform (57). In brief, reads were aligned to the GRCm38 (mm10) build of the mouse genome using TopHat2 (58) and FPKM values were computed with Cufflinks (59). The sequence data has been submitted to GEO repository and can be accessed using the accession number GSE84375. Differential gene expression analysis was carried out using DEseq2 (60) on the raw read counts based on release 82 of the Ensembl database. Finally, genes were selected according to a threshold of 2 for the fold change and 0.05 for the p-value.

### Analysis of glycolytic metabolites

Metabolites pertaining to glycolysis were analyzed with the assistance of Metabolomic Discoveries, Berlin, Germany. At the indicated time points post infection, BMDMs were washed with cold 0.9% NaCl and cells were collected in extraction buffer provided by Metabolomic Discoveries. Samples were snap frozen and sent to Metabolomic Discoveries. Derivatization and analysis of metabolites by a GC-MS 7890A mass spectrometer (Agilent, Santa Clara, USA) were carried out as described (45). Metabolites were identified in comparison to Metabolomic Discoveries database entries of authentic standards. The LC separation was performed using hydrophilic interaction chromatography with a ZIC-HILIC 3.5μm, 200A column (Merck Sequant, Umeå Sweden), operated by an Agilent 1290 UPLC system (Agilent, Santa Clara, USA). The LC mobile phase was A) 95% acetonitrile; 5% 10 mM ammonium acetate and B) 95% 10mM ammonium acetate; 5% acetonitrile with a gradient from 95 % A to 72 % A at 7 min, to 5% at 8 min, followed by 3 min wash with 5% A. The flow rate was 400μl/min, injection volume 1 μl. Mass spectrometry was performed using a 6540 QTOF/MS Detector and an AJS ESI source (Agilent, Santa Clara, USA). The measured metabolite intensities were normalized to internal standards.

### Glucose uptake assay

After 0, 0.5, 1 or 2 h post-infection, RPMI medium containing glucose was replaced with medium without glucose supplemented with 10 µM of the fluorescent glucose analogue 2-NBDG (Life Technologies). After 30 min of incubation at 37°C, cells were washed with PBS and resuspended in 1% formaldehyde for FACS analysis. FACS Canto (BD biosciences) flow cytometer was used for the acquisition of samples and Flowjo software (Flowjo LLC) was used for data analysis.

### Extracellular acidification rate measurement (Seahorse)

Extracellular acidification rate (ECAR) was analyzed using a XF96 Extracellular Flux Analyzer (Seahorse Bioscience). BMDMs were infected with *S*. Typhimurium with a MOI of 10 plated in non-buffered media. Measurements were obtained under basal conditions.

### Phagosomal β-galactosidase activity assay

To assess the β-galactosidase activity in phagolysomes, red fluorescent beads (Bangs Laboratories) were coated with 5-Dodecanoylaminofluorescein Di-β-D-Galactopyranoside (C_12_FDG, Life Technologies) for 60 min at 37°C in NaHCO_3_ pH 9.6 buffer. 100 beads per cell were added to BMDMs and incubated for 10 min at RT and 10 min at 37°C, followed by washings with RPMI to remove extracellular beads. After 0h, 0.5h, 1h or 2h, cells were washed with cold PBS and resuspended in 1% formaldehyde. Samples were acquired in a FACSCanto flow cytometer and Flowjo software. Mean fluorescence intensity (MFI) of C_12_FDG was normalized to the red MFI of the beads for every sample.

### Phagosomal proteolytic activity assay

To assess proteolytic activity in the phagolysosmes, red fluorescent beads were coated with green DQ-BSA (Life Technologies) dissolved in carbodiimide solution (25 mg/mL in PBS) for 30 min at RT. After washing, beads were resuspended in 0.1 M sodium tetraborate decahydrate solution (pH: 8.0 in ddH_2_O) and incubated over night at RT. Beads were then washed, resuspended in RPMI and added to BMDMs at 100beads/cell. After 10min incubation at RT and 10min incubation at 37°C, cells were washed to remove non-internalized beads. After 0, 0.5, 1 or 2h, cells were washed with cold PBS and resuspended in 1% formaldehyde. Samples were acquired using a BD FACSCanto flow cytometer and Flowjo software. DQ-BSA MFI was normalized to the red MFI of beads for every sample.

### Phagosomal acidification

Phagosome acidification was analyzed using the pH-sensitive fluorescent pHrodo Green conjugated *E. coli* Bioparticles (Life Technologies). These particles were first coated with the pH-insensitive dye Alexa Fluor-647 (Life Technologies) for 1 h at 37°C in 0.1M sodium tetra borate decahydrate (pH: 8.0 in ddH_2_O). Beads were then washed, resuspended in RPMI with 10 % FBS and added to the cells (100 bioparticles per sample). After 10 min incubation at RT and 10 min incubation at 37°C, cells were washed to remove non-internalized bioparticles. After 0h, 0.5h, 1h or 2h, cells were washed with cold PBS and resuspended in 1% formaldehyde. Fluorescent signal was analyzed using a FACSCanto flow cytometer and Flowjo software. pHrodo MFI was normalized to the Alexa-647 MFI for every sample.

### Antigen presentation assay

Treated BMDMs were incubated with Fc receptor blocking reagent TruStain fcX (Biolegend) for 5 min on ice and then incubated for 30 min on ice with an antibody specific for MHC class II or OVA_323-339_-MHC II complexes in PBS with 3% FBS solution. After washing, cells were resuspended in 1% formaldehyde in PBS and samples were acquired using a BD FACSCanto flow cytometer and analyzed using Flowjo software.

### OVA processing assay

Octadecyl C18 1µm magnetic beads (SiMAG, Chemicell) were coated with Alexa647-OVA (Life Technologies) in acetate buffer pH: 5.0 for 2 h at RT. After washing, beads were added to treated or infected BMDMs with a ratio of 300 beads per cell. After 24 h, cells were lysed, and Alexa647-OVA-coated magnetic beads were extracted as described above for the isolation of bead phagosomes. Isolated phagosomes were then resuspended in 1% formaldehyde (in PBS) and Alexa647-OVA signal was acquired and analyzed using a FACSCanto flow cytometer and Flowjo software respectively.

### Isolation of bead-containing phagosomes

BMDMs were incubated with Octadecyl C18 1 µm magnetic beads (SiMAG, Chemicell; 300 beads per cell) for 10 min at RT and for 10 min at 37°C. At each time point, cells were washed with PBS and Equilibration buffer (50 mM PIPES pH: 7.0, 50 mM MgCl_2_, 5 mM EGTA, 1 mM DTT, 10 µM cytochalasin and protease/phosphatase inhibitor cocktail) was added. Cells were incubated on ice for 20 min and samples were lysed (50 mM PIPES pH: 7.0, 50 mM MgCl_2_, 5 mM EGTA and 68mM sucrose). Cells were scrapped out and passed through a 26G needle at least 15 times. Bead-containing phagosomes were then separated from the lysate using a magnet.

### Isolation of *S*. Typhimurium-containing phagosomes

*S*. Typhimurium*-*containing phagosomes were isolated as described before (61). *S*. Typhimurium was grown in BHI broth until OD_600_:1 and then biotinylated with EZ-link NHS-Biotin reagent (Thermo Fisher Scientific). After washing, biotinylated bacteria were incubated with siMAG Streptavidin ferrofluid (Chemicell). Biotinylated *S*. Typhimurium bound to the Streptavidin ferrofluid was then separated using a magnet and bacteria were quantified using BHI agar plates. Subsequently, BMDMs were infected with biotinylated *S*. Typhimurium bound to the Streptavidin ferrofluid at an MOI of 10. At each time point, *S*. Typhimurium-containing phagosomes were isolated using equilibration and lysis buffer as described for bead-containing phagosome isolation.

### *In vitro* bacterial burden and ELISA

After 0 and 24h post-infection, BMDMs were lysed with 1% Triton X-100, 0.01% SDS in PBS. Several dilutions of the lysate were plated on BHI plates and incubated over night at 37°C. Next day, *S*. Typhimurium colony forming units (CFU) were enumerated. Supernatants were collected and analyzed for IL-6 and TNFα secretion using ELISA kit (R&D) according to the manufacturer’s instructions.

### Estimation of bacterial burden *in vivo*

Mice were infected with 100 CFU of *S*. Typhimuirum per mouse by i.v. injection. After 3 days of infection mice were euthanized according to current ethical protocols. Liver was isolated and homogenized using gentleMACS Dissociator (Miltenyi Biotec) in sterile PBS. Extracts of the homogenized livers were plated on BHI Agar plates. After 24 h incubation at 37°C, bacterial colonies were enumerated. Number of colonies was normalized to per gram of tissue. Mice were injected with 4-OHT intraperitoneally one day before the infection with *S*. Typhimurium and every day during the course of infection until the mice were sacrificed for analysis.

### Immunoblot analysis

BMDMs were lysed in RIPA buffer supplemented with 1X protease/phosphatase inhibitor cocktail (Thermo Fisher Scientific). Protein was estimated using Pierce® BCA Protein Assay Kit (Thermo Fisher Scientific) according to the manufacturer’s instructions. Equal amount of proteins was separated in 10% or 12% SDS-PAGE gels. Proteins were then transferred onto a PVDF membrane and probed with antibodies against Glut1 (sc-7903, Santa Cruz Biotechnology), Enolase (sc-31859, Santa Cruz Biotechnology), HKX-2 (sc-6521, Santa Cruz Biotechnology), v-ATPase A1 (V_0_ subunit, sc-28801, Santa Cruz Biotechnology), v-ATPase b1/2 (V_1_ subunit, sc-21209, Santa Cruz Biotechnology), H2-I/Abβ (sc-71201, Santa Cruz Biotechnology), phospho-p38 (#9216, Cell Signaling), p38 (#9212, Cell Signaling), phospho-p65 (#3033, Cell Signaling), p65 (#4764, Cell Signaling), HIF1α (NB100-105, Abcam), MondoA (SAB2104303, Sigma-Aldrich), calnexin (sc-11397, Santa Cruz Biotechnology) or β-actin (sc-47778, Santa Cruz Biotechnology). Either calnexin or β-actin was used as loading control. After incubation with secondary HRP-conjugated antibody (R&D) blots were developed using ECL reagent (GE Healthcare).

### Immunoprecipitation and Native PAGE

Bead phagosomes were isolated from 2-DG-pre-treated BMDMs as described above and were lysed with radio-immunoprecipitation assay (RIPA) buffer containing protease and phosphatase inhibitors. After preclearing the cell lysate with protein A/G agarose magnetic beads (#16-663, Millipore) for 1 h, beads were removed by placing the tube on a magnetic rack. The whole cell lysate (approximately 500 μg of protein) was incubated with 4 µg of an antibody against V_0_ subunit-*a* overnight at 4°C. A separate sample was incubated with IgG which served as a control. Protein A/G agarose beads were added again and incubated for an additional 1 h at room temperature. The immunoprecipitated proteins along with the agarose beads were collected by placing the tube on a magnetic rack. The collected beads were washed several times with RIPA buffer. The washed samples were mixed with SDS-PAGE sample loading buffer, boiled and resolved on a 10% SDS-polyacrylamide gel. V_1_ subunit B immunoprecipitated along with V_0_ was identified by Western blot analysis.

To perform Native-PAGE, equal amounts of total protein per sample were mixed with NativePAGE™ Sample Buffer (Life Technologies) and Triton X-100 (final concentration of 0.5%). Sample proteins were separated according to their masses on a 3.5 to 16% linear gradient acrylamide gel by electrophoresis. After separation, proteins were transferred to a PVDF membrane. Following blocking, membrane was incubated with antibodies directed against the cytosolic subunit of the v-ATPase (anti-vATPase-V_1_ subunit B) or against the membrane subunit of the v-ATPase (anti-vATPase-V_0_ subunit *a*).

### Statistical analysis

Statistical analysis was performed using Graphpad Prism software. Two-tailed Student’s *t*-test was conducted for most of the datasets unless specified otherwise to determine statistical significance. All data are represented as mean ± SEM as indicated. For all tests, p-values <0.05 was considered statistically significant (*p<0.05; **p<0.01; ***p<0.005).

## Author Contributions

Conceptualization: N.R., and S.G; Methodology: S.G., R.G., A.P., P.F., V.R.V., N.R., Investigation and Intellectual Input: S.G., J.F., R.G., M.W., A.P., G.C and N.R.; Writing Original Draft: S.G., and N.R.; Writing Review & Editing: P.F., V.R.V., G.C and N.R.; Funding Acquisition: N.R.; Resources: N.R., V.R.V: Supervision: V.R.V and N.R.

## Acknowledgements

This work was supported by funding to NR from Cologne Excellence Cluster on Cellular Stress Responses in Aging-Associated Diseases (CECAD; funded by the DFG within the Excellence Initiative by the German federal and state governments) Köln Fortune and grants from Deutsche Forschungsgemeinschaft (SFB 670) to NR. We thank Adam Antebi and Nina J. Hos for critically reading the manuscript.

## References

1. Ginhoux F, Schultze JL, Murray PJ, Ochando J, & Biswas SK (2016) New insights into the multidimensional concept of macrophage ontogeny, activation and function. Nat Immunol 17(1):34–40.

2. Robinson N, et al. (2012) Type I interferon induces necroptosis in macrophages during infection with Salmonella enterica serovar Typhimurium. Nature immunology 13(10):954–962.

3. Zhang DW, et al. (2009) RIP3, an energy metabolism regulator that switches TNF-induced cell death from apoptosis to necrosis. Science 325(5938):332–336.

4. Delmastro-Greenwood MM & Piganelli JD (2013) Changing the energy of an immune response. Am J Clin Exp Immunol 2(1):30–54.

5. Krawczyk CM, et al. (2010) Toll-like receptor-induced changes in glycolytic metabolism regulate dendritic cell activation. Blood 115(23):4742–4749.

6. Galvan-Pena S & O’Neill LA (2014) Metabolic reprograming in macrophage polarization. Front Immunol 5:420.

7. Rodriguez-Prados JC, et al. (2010) Substrate Fate in Activated Macrophages: A Comparison between Innate, Classic, and Alternative Activation. J Immunol 185(1):605–614.

8. Huang SCC, et al. (2014) Cell-intrinsic lysosomal lipolysis is essential for alternative activation of macrophages. Nat Immunol 15(9):846–855.

9. Kornberg MD, et al. (2018) Dimethyl fumarate targets GAPDH and aerobic glycolysis to modulate immunity. Science 360(6387):449–453.

10. Mills EL, et al. (2018) Itaconate is an anti-inflammatory metabolite that activates Nrf2 via alkylation of KEAP1. Nature 556(7699):113–117.

11. Tannahill GM, et al. (2013) Succinate is an inflammatory signal that induces IL-1 beta through HIF-1 alpha. Nature 496(7444):238-+.

12. Eisele NA, et al. (2013) Salmonella Require the Fatty Acid Regulator PPAR delta for the Establishment of a Metabolic Environment Essential for Long-Term Persistence. Cell Host Microbe 14(2):171–182.

13. Bowden SD, Rowley G, Hinton JC, & Thompson A (2009) Glucose and glycolysis are required for the successful infection of macrophages and mice by Salmonella enterica serovar typhimurium. Infect Immun 77(7):3117–3126.

14. Ganesan R, et al. (2017) Salmonella Typhimurium disrupts Sirt1/AMPK checkpoint control of mTOR to impair autophagy. PLoS pathogens 13(2):e1006227.

15. Fischer J, et al. (2019) Leptin signaling impairs macrophage defenses against Salmonella Typhimurium. Proceedings of the National Academy of Sciences of the United States of America 116(33):16551–16560.

16. Galic S, Sachithanandan N, Kay TW, & Steinberg GR (2014) Suppressor of cytokine signalling (SOCS) proteins as guardians of inflammatory responses critical for regulating insulin sensitivity. Biochem J 461(2):177–188.

17. Weng LP, Smith WM, Brown JL, & Eng C (2001) PTEN inhibits insulin-stimulated MEK/MAPK activation and cell growth by blocking IRS-1 phosphorylation and IRS-1/Grb-2/Sos complex formation in a breast cancer model. Hum Mol Genet 10(6):605–616.

18. Kelly B & O’Neill LA (2015) Metabolic reprogramming in macrophages and dendritic cells in innate immunity. Cell Res 25(7):771–784.

19. Freemerman AJ, et al. (2014) Metabolic reprogramming of macrophages: glucose transporter 1 (GLUT1)-mediated glucose metabolism drives a proinflammatory phenotype. J Biol Chem 289(11):7884–7896.

20. Luhrmann A & Haas A (2000) A method to purify bacteria-containing phagosomes from infected macrophages. Methods Cell Sci 22(4):329–341.

21. Mauer J, et al. (2010) Myeloid cell-restricted insulin receptor deficiency protects against obesity-induced inflammation and systemic insulin resistance. Plos Genet 6(5):e1000938.

22. Doughty CA, et al. (2006) Antigen receptor-mediated changes in glucose metabolism in B lymphocytes: role of phosphatidylinositol 3-kinase signaling in the glycolytic control of growth. Blood 107(11):4458–4465.

23. Kohn AD, et al. (1998) Construction and characterization of a conditionally active version of the serine/threonine kinase Akt. J Biol Chem 273(19):11937–11943.

24. Lennon-Dumenil AM, et al. (2002) Analysis of protease activity in live antigen-presenting cells shows regulation of the phagosomal proteolytic contents during dendritic cell activation. J Exp Med 196(4):529–540.

25. Forgac M (2007) Vacuolar ATPases: rotary proton pumps in physiology and pathophysiology. Nat Rev Mol Cell Biol 8(11):917–929.

26. Kane PM (1995) Disassembly and reassembly of the yeast vacuolar H(+)-ATPase in vivo. J Biol Chem 270(28):17025–17032.

27. Lu M, Sautin YY, Holliday LS, & Gluck SL (2004) The glycolytic enzyme aldolase mediates assembly, expression, and activity of vacuolar H+-ATPase. The Journal of biological chemistry 279(10):8732–8739.

28. Su Y, Zhou A, Al-Lamki RS, & Karet FE (2003) The a-subunit of the V-type H+-ATPase interacts with phosphofructokinase-1 in humans. The Journal of biological chemistry 278(22):20013–20018.

29. O’Neill LA & Pearce EJ (2016) Immunometabolism governs dendritic cell and macrophage function. J Exp Med 213(1):15–23.

30. Mehrotra P, et al. (2014) Pathogenicity of Mycobacterium tuberculosis Is Expressed by Regulating Metabolic Thresholds of the Host Macrophage. Plos Pathog 10(7).

31. Hernandez LD, Pypaert M, Flavell RA, & Galan JE (2003) A Salmonella protein causes macrophage cell death by inducing autophagy. J Cell Biol 163(5):1123–1131.

32. Layton AN, Brown PJ, & Galyov EE (2005) The Salmonella translocated effector SopA is targeted to the mitochondria of infected cells. J Bacteriol 187(10):3565–3571.

33. Hos NJ, et al. (2017) Type I interferon enhances necroptosis of Salmonella Typhimurium-infected macrophages by impairing antioxidative stress responses. The Journal of cell biology 216(12):4107–4121.

34. Liu XY, Lu R, Xia YL, & Sun J (2010) Global analysis of the eukaryotic pathways and networks regulated by Salmonella typhimurium in mouse intestinal infection in vivo. Bmc Genomics 11.

35. Antunes LCM, et al. (2011) Impact of Salmonella Infection on Host Hormone Metabolism Revealed by Metabolomics. Infect Immun 79(4):1759–1769.

36. Robinson N, et al. (2012) Type I interferon induces necroptosis in macrophages during infection with Salmonella enterica serovar Typhimurium. Nat Immunol 13(10):954–962.

37. Everts B, et al. (2014) TLR-driven early glycolytic reprogramming via the kinases TBK1-IKK epsilon supports the anabolic demands of dendritic cell activation. Nat Immunol 15(4):323-+.

38. Lambeth JD (2004) NOX enzymes and the biology of reactive oxygen. Nat Rev Immunol 4(3):181–189.

39. Sanman LE, et al. (2016) Disruption of glycolytic flux is a signal for inflammasome signaling and pyroptotic cell death. Elife 5:e13663.

40. Buchmeier NA & Heffron F (1991) Inhibition of Macrophage Phagosome-Lysosome Fusion by Salmonella-Typhimurium. Infect Immun 59(7):2232–2238.

41. Oh YK, et al. (1996) Rapid and complete fusion of macrophage lysosomes with phagosomes containing Salmonella typhimurium. Infect Immun 64(9):3877–3883.

42. Mills SD & Finlay BB (1998) Isolation and characterization of Salmonella typhimurium and Yersinia pseudotuberculosis-containing phagosomes from infected mouse macrophages: Y-pseudotuberculosis traffics to terminal lysosomes where they are degraded. Eur J Cell Biol 77(1):35–47.

43. Coombes BK, Brown NF, Valdez Y, Brumell JH, & Finlay BB (2004) Expression and secretion of Salmonella pathogenicity island-2 virulence genes in response to acidification exhibit differential requirements of a functional type III secretion apparatus and SsaL. Journal of Biological Chemistry 279(48):49804–49815.

44. Albaghdadi H, Robinson N, Finlay B, Krishnan L, & Sad S (2009) Selectively reduced intracellular proliferation of Salmonella enterica serovar typhimurium within APCs limits antigen presentation and development of a rapid CD8 T cell response. J Immunol 183(6):3778–3787.

45. Lisec J, Schauer N, Kopka J, Willmitzer L, & Fernie AR (2006) Gas chromatography mass spectrometry-based metabolite profiling in plants. Nat Protoc 1(1):387–396.

46. Vieira OV, Botelho RJ, & Grinstein S (2002) Phagosome maturation: aging gracefully. The Biochemical journal 366(Pt 3):689–704.

47. West AP, et al. (2011) TLR signalling augments macrophage bactericidal activity through mitochondrial ROS. Nature 472(7344):476–480.

48. Chakravortty D & Hensel M (2003) Inducible nitric oxide synthase and control of intracellular bacterial pathogens. Microbes Infect 5(7):621–627.

49. Sturgillkoszycki S (1994) Lack of Acidification in Mycobacterium Phagosomes Produced by Exclusion of the Vesicular Proton-Atpase (Vol 263, Pg 678, 1994). Science 263(5152):1359–1359.

50. Nordenfelt P, Grinstein S, Bjorck L, & Tapper H (2012) V-ATPase-mediated phagosomal acidification is impaired by Streptococcus pyogenes through Mga-regulated surface proteins. Microbes Infect 14(14):1319–1329.

51. Kohio HP & Adamson AL (2013) Glycolytic control of vacuolar-type ATPase activity: A mechanism to regulate influenza viral infection. Virology 444(1-2):301-309.

52. Havula E, et al. (2013) Mondo/ChREBP-Mlx-Regulated Transcriptional Network Is Essential for Dietary Sugar Tolerance in Drosophila. Plos Genet 9(4).

53. Chang CH, et al. (2013) Posttranscriptional control of T cell effector function by aerobic glycolysis. Cell 153(6):1239–1251.

54. Schindler A & Foley E (2013) Hexokinase 1 blocks apoptotic signals at the mitochondria. Cell Signal 25(12):2685–2692.

55. Roman-Garcia P, et al. (2014) Vitamin B(1)(2)-dependent taurine synthesis regulates growth and bone mass. J Clin Invest 124(7):2988–3002.

56. Xia J, Sinelnikov IV, Han B, & Wishart DS (2015) MetaboAnalyst 3.0--making metabolomics more meaningful. Nucleic Acids Res 43(W1):W251-257.

57. Wagle P, Nikolic M, & Frommolt P (2015) QuickNGS elevates Next-Generation Sequencing data analysis to a new level of automation. Bmc Genomics 16:487.

58. Kim D, et al. (2013) TopHat2: accurate alignment of transcriptomes in the presence of insertions, deletions and gene fusions. Genome Biol 14(4):R36.

59. Trapnell C, et al. (2010) Transcript assembly and quantification by RNA-Seq reveals unannotated transcripts and isoform switching during cell differentiation. Nat Biotechnol 28(5):511–515.

60. Love MI, Huber W, & Anders S (2014) Moderated estimation of fold change and dispersion for RNA-seq data with DESeq2. Genome Biol 15(12):550.

61. Gutierrez S, Wolke M, Plum G, & Robinson N (2017) Isolation of Salmonella typhimurium-containing Phagosomes from Macrophages. Journal of visualized experiments : JoVE (128).

